# DeepGEEP: Data-Driven Prediction of Bacterial Biofilm Gene Expression Profiles

**DOI:** 10.1101/2023.08.30.555510

**Authors:** Hamidreza Arjmandi, Christophe Corre, Hamidreza Jahangir, Adam Noel

## Abstract

Predicting the gene expression profile (GEEP) of bacterial biofilms in response to spatial, temporal, and concentration profiles of stimulus molecules holds significant potential across microbiology, biotechnology, and synthetic biology domains. However, the resource and time-intensive nature of experiments within Petri dishes presents significant challenges. Data-driven methods offer a promising avenue to replace or reduce such experiments given sufficient data. Through well-crafted data generation techniques, the data scarcity issue can be effectively addressed. In this paper, an innovative methodology is presented for generating GEEP data over a Petri dish that results from a specific chemical stimulus release profile. A twodimensional convolutional neural network (2D-CNN) architecture is subsequently introduced to leverage the synthesized dataset to predict GEEP variations across bacterial biofilms within the Petri dish. The approach, coined DeepGEEP, is applied to data generated by a particle-based simulator (PBS) to enable a flexible evaluation of its efficacy. The proposed method attains a significant level of accuracy in comparison to established benchmark models such as Linear SVM, Radial Basis Function SVM, Decision Tree, and Random Forest.

## 1. Introduction

**T**HE responses of cells within living organisms to diverse stimuli underscore the remarkable complexity of biological systems. These responses invariably entail the modulation of gene expression triggered by the corresponding stimulus. Gene expression refers to the process by which information encoded in a gene is used to synthesize a functional gene product, such as a protein or RNA molecule. It is the fundamental mechanism by which cells respond to external stimuli and adapt to their environment [1].

Gene expression can be detected and visualized in a Petri dish experiment typically using various molecular biology techniques or reporter systems. For example, chemically inducible promoters can be engineered to drive the expression and production of fluorescent proteins, such as Green Fluorescent Protein (GFP), that emit light when exposed to specific wavelengths. By fusing a promoter upstream of the GFP gene, we can track and visualize gene expression under a fluorescence microscope or even under the naked eye in specific conditions.

### A. Motivation

Bacteria are relatively simple living organisms but understanding their various gene functionalities could have a huge impact on our health and the environment [2]. Bacteria grow in biofilms as communities that adhere to surfaces. Therefore, bacteria culture growing in a Petri dish is a very common environment for *in vivo* experiments.

Predicting the GFP expression profile (GEEP) of a bacteria biofilm over a Petri dish based on the chemical stimulus release profile (i.e., space, time, and concentration) has several potential applications. GEEP prediction can help researchers understand how biofilms respond to different chemical stimulus release profiles, aiding in the development of strategies to control or enhance biofilm formation in various natural and engineered environments [3]. Also, profile prediction can help to identify novel targets for antimicrobial therapies and develop strategies to disrupt or prevent biofilm formation [4], [5]. Understanding how biofilm behavior changes in response to varying stimulus release profiles can enable us to optimize bioremediation processes and improve pollutant removal efficiency [6].

In biotechnology and industrial applications, biofilms are used for various purposes, such as biofuel production [7]. Predicting the GEEP can guide the design and optimization of bioprocesses involving biofilm formation [8]. Also, engineers and researchers working in synthetic biology can use the predicted GEEPs to design and engineer biofilms with specific behaviors or responses to different environmental stimuli. This could lead to the development of biosensors or biodevices that respond to changing conditions [9].

### B. Different Approaches for GEEP Prediction

Conducting GEEP experiments within Petri dishes can be both expensive and time-consuming [10]. Attempting to replicate these experiments across a broad spectrum of situations to encompass all potential scenarios often becomes infeasible and hinders the exploration of many biophysical hypotheses. Also, the outcomes of these experiments are susceptible to human errors, which can significantly impact the accuracy of results. Additionally, the extensive execution of such experiments contributes to environmental pollution, raising significant ecological concerns. Furthermore, personnel safety when conducting numerous experiments is another concern.

The drawbacks of experimental methodologies have paved the way for the adoption of computational modeling approaches, as highlighted in [11]. These computational strategies are broadly categorized into two distinct paradigms: traditional computational parametric models and data-driven models.

Researchers have increasingly used computational parametric models employing partial differential equations (PDEs) to unravel the intricate processes governing signaling within bacterial cultures, e.g., quorum sensing. This line of inquiry aims to elucidate the spatio-temporal distribution of signaling molecules and its consequential impact on gene expression [12]–[15]. In [19], a biophysical model is introduced to predict the time-dependent concentration distribution of N-acyl homoserine lactone (AHL) within an agarose matrix. Authors in [20] conducted an experiment where a bacterial strain was embedded in a short agar lane. An exogenous autoinducer was introduced at one end of the lane to track the expression of a quorum sensing (QS) reporter over time and space, enabling the measurement of autoinducer diffusion effects along the lane. Furthermore, a droplet-based bacterial communication system was devised in [21]. This system employed *Escherichia coli* (*E. coli*) bacteria enclosed within water-in-oil emulsion droplets of significant populations. This work demonstrated the potential of this system for genetically-programmed pattern formation and distributed computing.

Characterization of diffusion sensing by individual cells [16], the clustering patterns of bacterial aggregates within a biofilm matrix [17], the phenomena of collective sensing, and the adaptability of bacterial cultures to heterogeneous environments [18] are also facets of bacterial culture behavior that have received attention from a computational perspective. However, conventional parametric methods suffer from significant limitations. The accurate estimation of parameters within these experimental setups is not readily attainable through a limited number of trials. Moreover, their adaptability is constrained, and their ability to obtain insights from experience is limited.

Data-driven approaches might initially appear to contradict the very essence of our objectives within this paper. One might assume that such methodologies necessitate an extensive array of experiments to generate the requisite data. Nonetheless, through a thoughtfully designed data-driven approach, we can effectively tackle this challenge by leveraging our existing comprehension of the system. By thoughtfully defining features and data samples based on our knowledge of the underlying processes, we can generate an extensive dataset from a single experiment, thus mitigating the aforementioned concern. The utilization of data-driven methodologies to extract features from biofim images is increasing, such as individual cell geometry [23], and the classification of cells in comparison to other substances [24].

In this paper, we introduce a novel methodology for generating data, tailored to the data-driven prediction of gene expression profiles (DeepGEEP) within bacterial biofilms on a Petri dish in response to a specific chemical stimulus release profile. This approach is based on the notion that different cell colonies on identical plates have different distances from the release area points (sources) and Petri dish boundaries. We harness these attributes within the data generation technique to avoid the need to teach the machine to extract these features from a multitude of images corresponding to numerous experiments. Subsequently, we propose the utilization of a two-dimensional convolutional neural network (2D-CNN) architecture, which harnesses the generated dataset to predict the GEEP within the bacterial biofilm across the Petri dish.

### C. Contributions

In this study, we produce our data using a particle-based simulator (PBS). This simulator traces the movement of diffusing molecules within the Petri dish environment, which enables us to assess the viability of our approach. The PBS includes diffusion within the biofilm and agar, boundary conditions, and a simplified gene expression process. Our research introduces several notable contributions:

- We present DeepGEEP as a deep learning strategy aimed at predicting Green Fluorescent Protein (GFP) expression profiles across the Petri dish, offering a potential reduction in the need for numerous costly and time-consuming experiments. This approach serves to address the limitations associated with computational parametric methods.
- We propose a novel approach to generate data from a limited number of simulations (experiments) to train the deep learning model. This method involves sampling and segmenting the experimental (simulation) environment, enabling us to produce a diverse array of data samples from each experiment (simulation).
- We introduce a specific 2D-CNN architecture that achieves a notably high level of accuracy when compared to benchmark models such as Linear support vector machine (SVM), Radial Basis Function SVM, Decision Tree, and Random Forest.

## II. System Model and Problem Statement

In this section, we provide an overview of the fundamental model involving an agar Petri dish, which serves as the environment for bacterial cultivation, and the controlled release of a chemical stimulus.

We assume a cylindrical Petri dish with a radius of *r*_*p*_ m filled with agar to a height of *h*_*a*_ m. We employ cylindrical coordinates with the origin situated at the center of the top of the agar. The coordinates (*r, θ, z*) correspond to the radial, azimuthal, and axial dimensions, respectively.

The starting bacterial culture, diluted with water, has a concentration denoted as *C*_*b*_ cel·m^−31^. This concentration can be practically quantifiable through optical spectroscopy. A specified volume of *V*_*b*_ m^3^ from this bacterial source is uniformly distributed over the entirety of the top agar surface. We consider the dilution of signaling molecules by mixing them with water, resulting in a concentration denoted as *C*_*m*_ mol·m^−3^. A droplet of volume *V*_*d*_ m^−3^ of this solution is deposited at a specified point (*r*_0_, *θ*_0_, *z*_0_) on the agar surface, forming an initial area of *A*_*d*_ m^2^. As time progresses, the droplet, carrying the signaling molecules, infiltrates the diffusible agar medium and undergoes a reduction in size, resulting in a time-varying area denoted as *A*_*d*_(*t*). It is important to note that the height of the droplet, denoted as *h*_*d*_, is assumed fixed during the soaking which is determined by the ratio of droplet volume to the initial area, calculated as *V*_*d*_*/A*_*d*_(0). The signaling molecules diffuse through the agar matrix and may interact with cells present on the surface through their diffusive movement. The interaction can stimulate the cells and thus generate a response, e.g., express a specific gene and produce the corresponding protein.

After dropping the droplet onto the plate, the plate is placed in an incubator (37°C) to enhance the growth of the bacteria and to form a biofilm. The cells’ response to the stimulus may vary over the plate depending on several factors including the concentration of received molecules, cell-to-cell variation, and cell-to-cell communication. Given that the bacteria have been engineered to produce GFP in response to the signalling molecules, then we can observe a green pattern over the Petri dish when exposed to blue light. The goal is to predict the cells’ response pattern (i.e., gene expression pattern) over the Petri dish given the release area of the droplet. Fig. 1 depicts the schematic of the system model.

**Fig. 1:**
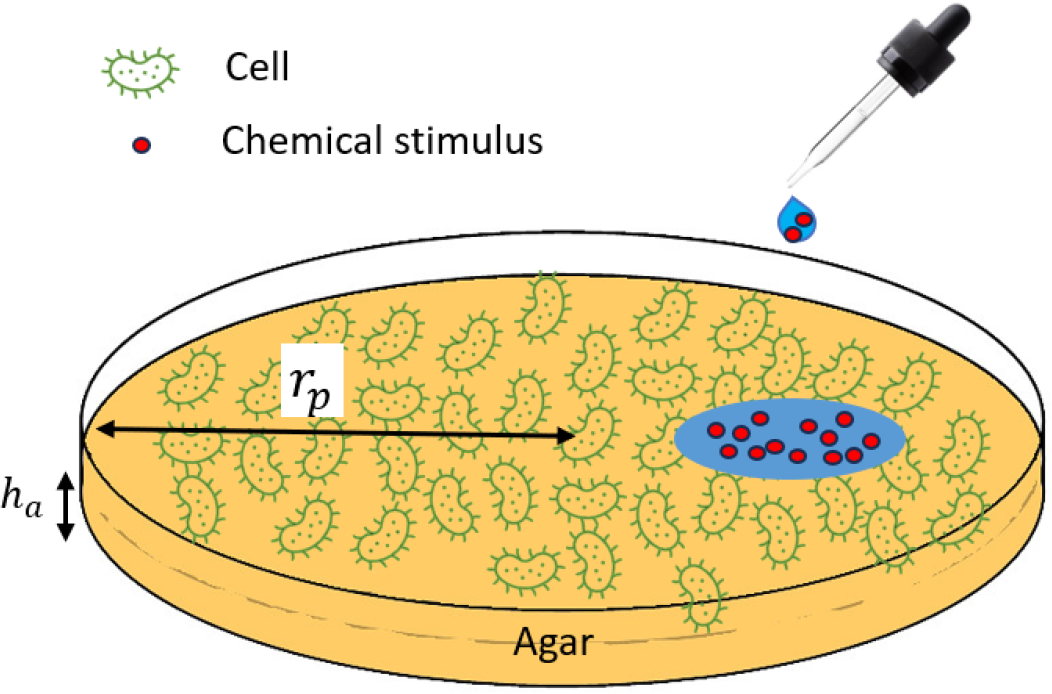
Schematic of the system model. A droplet carrying the chemical stimulus is deposited at a specific point on the agar surface that is hosting the cells (the cells and molecules are not depicted to scale).

### Example

An experiment conducted in molecular microbiology laboratory studies focuses on the *E. coli* bacteria and Isopropyl *β*-D-1-thiogalactopyranoside (IPTG) as the signaling molecule. In *E. coli* research, the gene responsible for encoding GFP is frequently controlled by an inducible promoter (e.g., the lac promoter) that responds to IPTG via the interaction of IPTG with the trancripsional repressor protein LacI. This process enables researchers to regulate GFP expression by introducing IPTG. The pairing of an IPTG-responsive promoter with GFP forms a reporter gene construct used for visualizing gene expression within *E. coli*.

## III. Data Generation

To generate the data, we employ a particle-based simulator. Leveraging the PBS enables us to modify parameters and systematically introduce, adjust, or modify the relevant processes as needed to investigate the feasibility of our approach for future experimental data. This strategy is notably efficient in terms of both time and cost.

### A. Particle-based Simulator

We begin by elaborating on the processes involved in *E. coli* GFP expression in response to releasing IPTG over the agar surface. Subsequently, we outline how the PBS effectively replicates each of these key processes involved in the experiment.

*1) Bacteria Spreading:* As previously indicated, a solution containing bacteria (bacteria diluted in sterile water) with a concentration denoted as *C*_*b*_ is uniformly spread across the entire surface.

*2) Droplet soaking:* Following the application of the bacteria solution, a droplet of IPTG solution is introduced. This droplet, with a volume of *V*_*d*_ m^−3^ and the concentration *C*_*m*_, is dropped at coordinates (*r*_0_, *θ*_0_, *z*_0_) on the agar surface. This leads to the formation of a circular region with an area of *A*_*d*_ m^2^ and an initial height of *h*_*d*_.

Given the soaking rate of *k*_*s*_ m s^−1^ for the droplet into agar, the temporal evolution of the droplet’s area is calculated as [22]

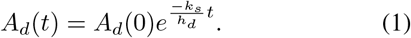

This soaking process serves as the molecule source in the PBS. To simulate this process within a given interval [*t*, Δ*t*], we account for the quantity of molecules that permeate the agar due to the soaking rate during this timeframe. Over this period, the droplet’s area reduces from *A*_*d*_(*t*) to *A*_*d*_(*t*+Δ*t*). Consequently, the corresponding change in volume is computed as *h*_*d*_(*A*_*d*_(*t*) − *A*_*d*_(*t* +Δ*t*)), which equates to *C*_*d*_*h*_0_(*A*_*d*_(*t*) − *A*_*d*_(*t* + Δ*t*)) molecules. It is assumed that this quantity of molecules infiltrates uniformly across the area *A*_*d*_(*t*) of the droplet into the agar.

*3) Diffusion in agar:* Molecules residing within the agar are assumed to undergo free diffusion. At each time increment, the positions of information molecules are adjusted in accordance with stochastic Brownian motion. This occurs independently for each molecule. The displacement of a molecule along each spatial dimension in Δ*t* seconds adheres to a Gaussian distribution with a mean of zero and a variance of 2*D*_*a*_Δ*t* in which *D*_*a*_ is the diffusion coefficient of IPTG in the agar. If a molecule’s trajectory during this temporal span results in it colliding with the plate’s boundaries, then it is redirected back within the agar. Similarly, should a molecule intersect the surface, it may adhere to bacteria with some probability, or it can be reflected back.

*4) IPTG consumption by bacteria:* Once spread across the surface, bacteria grow at a rate that is accelerated within an incubator with elevated temperatures. The interaction between IPTG and bacteria on the surface depends on the population of bacteria present. For the sake of simplicity, we consider an irreversible reaction to model molecules (M) consumption by cells (C) that sequesters the molecule from the environment:

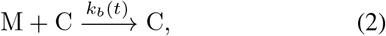

where *k*_*b*_(*t*) is the consumption rate of cell population area over the surface at time *t*. As cell growth progresses across the surface, the rate of consumption amplifies with time. To this end, we consider that the consumption rate increases linearly with the cell population. This assumption is described by the following equation:

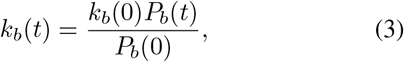

where *P*_*b*_(*t*) represents the cell population at time *t*.

*5) Bacteria growth over the plate:* Bacteria undergo a proliferation process and develop a biofilm structure. During the growth phase, the *E. coli* cell population doubles very 20 minutes. However, this exponential proliferation eventually halts due to spatial and nutritional constraints. Based on experimental observations [22], we assume that bacterial growth ceases after *H*_*b*_ hours which is marked by an absence of discernible bacterial boundary expansion. Consequently, by considering the initial consumption rate *k*_*b*_(0), we can infer the time-dependent consumption rate during the growth stage as follows:

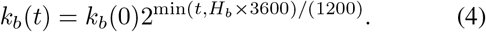

In cases where a molecule hits the bacterial surface, it has a chance of being consumed during a time interval Δ*t* seconds with a probability of approximately 1− exp (− *k*_*b*_(*t*)Δ*t*) ∼ *k*_*b*_(*t*)Δ*t*.

*6) Biofilm formation and diffusion in biofilm:* Another critical aspect of this experiment pertains to the moment of biofilm formation. At this time, molecules that hit the biofilm surface have the opportunity to infiltrate it, leveraging the water channels within to facilitate their diffusion [25]. We postulate that the biofilm forms after a duration of *T*_*bf*_ hours. Following this threshold time, molecules colliding with the surface can perfectly access the biofilm’s water channels, resulting in significantly accelerated movement compared to diffusion within the agar medium. While the actual structure of these water channels is intricate [25], we simplify its representation as a form of free-surface diffusion, characterized by an effective diffusion coefficient *D*_*b*_ greater than that of the agar diffusion coefficient *D*_*a*_. Assuming that molecules hitting the surface after *T*_*bf*_ enter the biofilm, we can describe their behavior during a time increment Δ*t*. During this interval, a molecule may either be consumed by bacteria with a probability of 1− exp (− *k*_*b*_Δ*t*), or it can engage in random movement along the x and y directions, characterized by Gaussian-distributed random variables with zero mean and a variance of 2*D*_*b*_Δ*t*.

In the event that a molecule is consumed/sequestered by a bacterium, it becomes permanently fixed at that location. This process results in the gradual consumption of nearly all molecules across the plate over time.

*7) Gene Expression:* Our experimental results [22] exhibit that bacteria respond to a specific range of concentration of IPTG. When the IPTG concentration surpasses a certain threshold, the bacteria might fail to exhibit a response or experience lethality. Conversely, concentrations below this range do not elicit gene expression. Based on this observation, we establish a threshold range [*C*_*L*_, *C*_*U*_], signifying the concentration levels within which the bacteria exhibit GFP expression in response to GFP.

## IV. DeepGEEP Design

In this section, we present our proposed approach to generate training data that enables GEEP using a limited number of experiments. Also, we provide the designed 2D-CNN structure corresponding with the generated training data.

### A. Training Data Generation

One can define different training data models from lab experiments. An intuitive method involves creating many distinct droplet areas and concentrations as training input data. These data take the form of 2D images, encapsulating the relevant information. In parallel, the resultant GFP patterns corresponding to each input image serve as the training output data. As a result, each experiment provides one sample for training. However, the execution of numerous time-consuming and expensive lab experiments poses a significant challenge and requires a resolution.

We introduce a novel approach that enables the resourceful generation of data using a limited number of experiments. The premise of this approach rests upon the notion that different cell colonies on identical plates have different distance profiles from the release area points and boundaries. These attributes can be preprocessed and harnessed within the data generation for the machine learning framework, avoiding the need to teach the machine to extract these features from a multitude of images corresponding to numerous experiments.

Consider the smallest square that completely encloses the circular plate. Now, picture this square being divided both vertically and horizontally into *N*_*x*_ equal parts. This division results in a matrix with dimensions *N*_*x*_ × *N*_*x*_. In this matrix, each element, represented by (*j, l*), indexes for a small square centered at a specific point (*X*_*j*_, *Y*_*l*_). Fig. 2 illustrates an 8 × 8 matrix that partitions the plate into 64 squares, with 32 of them fully located within the circle.

**Fig. 2:**
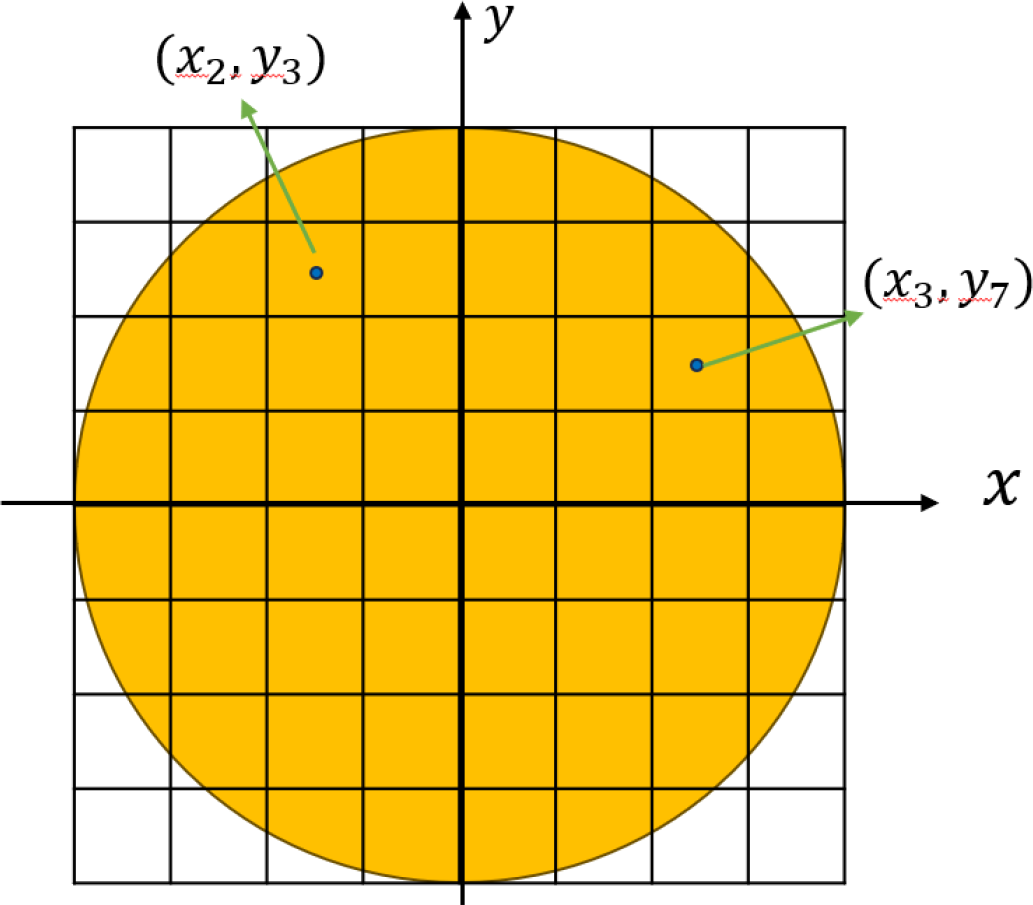
Partitioning the plate using a 8 × 8 matrix.

We denote the total number of square pieces (i.e., samples) fully *inside* the circular plate as *N*_*s*_ and each one will represent a group of bacteria. Each sample *I* ∈ 1, …, *N*_*s*_ is associated with a specific element in our matrix, let’s call it (*j*(*i*), *l*(*i*)). The center of matrix element (*j*(*i*), *l*(*i*)) is located at (*X*_*j*(*i*)_, *Y*_*l*(*i*)_) over the surface of the plate.

To make things simple, we label each sample with a value for GFP expression, denoted as *GFP* (*i*). This value can either be 0 or 1. When *GFP* (*i*) = 1, it means that the group of bacteria at (*X*_*j*(*i*)_, *Y*_*l*(*i*)_, 0) has triggered the production of GFP. In the PBS, this indicates that the number of IPTG molecules absorbed within this square is within the interval [*C*_*L*_, *C*_*U*_]. From the perspective of the experiments, this can be interpreted to mean that the observed fluorescence level is brighter.

The position of each sample *i* ∈ 1 …, *N*_*s*_, is incorporated as a component of our training input data. For each individual sample *i*, we define a grid matrix, *ℳ*_1_. In this matrix, every element is initially set to zero except for the element (*j*(*i*), *l*(*i*)), which corresponds exactly to the location of that particular sample, namely (*X*_*j*(*i*)_, *Y*_*l*(*i*)_), i.e.,

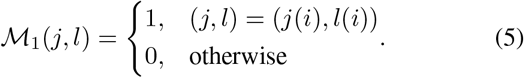

The distance between each point within the system and the sample point holds substantial informative value for the learning process due to the impact of diffusion. Thus, we establish another grid matrix named ℳ_2_. Within this matrix, each element (*j, l*) is calculated as the Euclidean distance between (*X*_*j*_, *Y*_*l*_) and the location of the sample at (*X*_*j*(*i*)_, *Y*_*l*(*i*)_), then normalized by a factor of 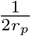, i.e., we have

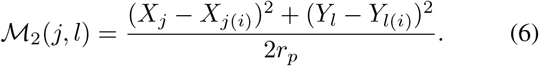

Another factor influencing the GFP expression (concentration) for a given sample is its positioning relative to the points along the side boundary of the plate. To account for this aspect, we introduce the grid matrix ℳ_3_ for each sample *i*. In this matrix, every element (*j, l*) located along the plate’s border is determined by the square of the Euclidean distance between (*X*_*j*_, *Y*_*l*_) and the position of the sample at (*X*_*j*(*i*)_, *Y*_*l*(*i*)_). This value is then normalized by a factor of 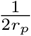. In other words, we calculate this distance-related effect for every element at the plate’s border, considering its relationship to the sample’s location, i.e.,

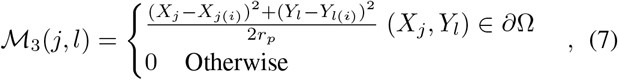

in which *∂*Ω represents the Petri dish boundary. Obviously, boundary points that are further from the sample have less impact.

Another aspect that the input data should account for is the region where the stimulus molecules are released, as well as the quantity of molecules released. One challenge that might arise is the impact of molecule release time. However, we can simplify the situation by ignoring the molecule release time during the soaking. This assumption stems from the fact that molecules move independently, and we are observing the system at a steady state. Consequently, the timing of molecule release becomes irrelevant; what matters is where molecules are released.

The total number of molecules released within each square piece of the designated release area is determined using the PBS capturing the droplet soaking mechanism outlined in subsection III-A. This allows the PBS to effectively estimate the total molecules released within each square area. As a result, a portion of the input data is represented by a grid matrix, ℳ_4_, where each element (*j, l*) denotes the accumulated number of molecules released within its corresponding region denoted by *N*_Rel_(*j, l*). It is evident that the maximum molecule count would be at the droplet’s center, while the count decreases as we move towards the initial border of the droplet. In all other areas, the molecule count is zero. Therefore, we have

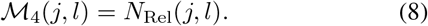

It is important to recognize that ℳ_4_(*j, l*) for a sample does not provide sufficient information to differentiate between the samples. This is because it remains the same for all samples within a single experiment or simulation. To enhance the efficiency of the training process, we introduce the grid matrix *M*_5_ whose element (*j, l*) within the release area is determined by the square of the Euclidean distance between (*X*_*j*_, *Y*_*l*_) andthe location of the sample at (*X*_*j*(*i*)_, *Y*_*l*(*i*)_), i.e.,

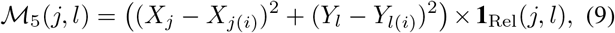

in which **1**_Rel_(*j, l*) is a function that indicates whether the (*j, l*) element corresponds to a release point. We chose the square of the Euclidean distance because the peak concentration at a point in an unbounded environment is inversely proportional to the square of the distance between the release and observation points [26].

### B. 2D-CNN Structure

In our model, the input data resembles an image of dimensions *N*_*x*_ × *N*_*y*_ with five channels. This suggests that we use a 2D-CNN structure for this type of data. Fig. 3 presents an overview of the architecture of the 2D-CNN devised for our proposed data structure to identify the gene expression pattern. Given *N*_sim_ simulation senarios, the input dataset takes the form of *N*_sim_×*N*_*s*_ samples, each with dimensions *N*_*x*_×*N*_*y*_ ×5, while the output consists of binary classifications denoted as {0, 1};.

**Fig. 3:**
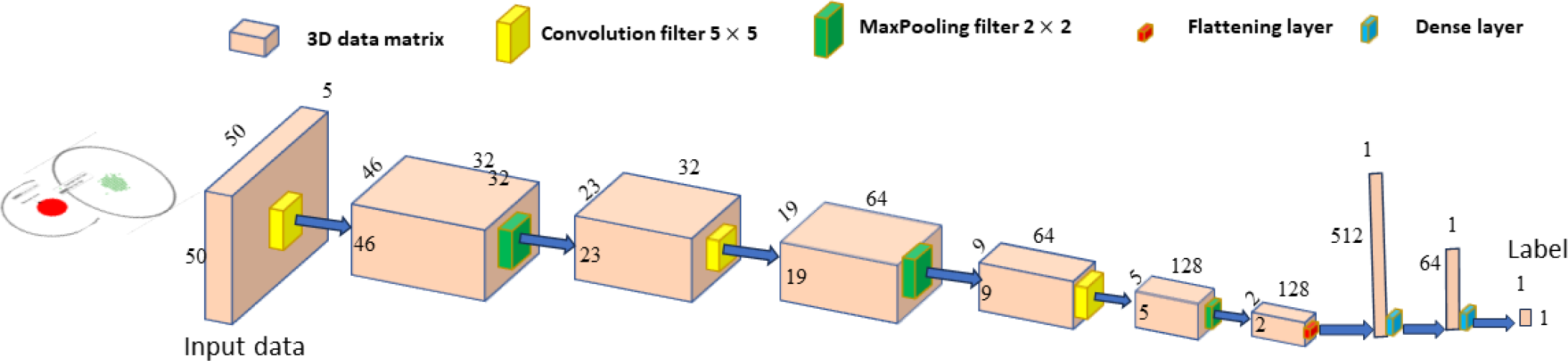
Schematic structure of 2D-CNN

The designed architecture of the 2D-CNN is as follows: the first layer is a 2D convolutional layer with 32 filters using the ReLU activation function. The input shape for this layer is (*N*_*x*_, *N*_*y*_, 5), which corresponds to the size of each input sample. After the first convolutional layer, a 2D MaxPooling layer with a pool size of (2, 2) is applied. MaxPooling reduces the spatial dimensions of the feature maps, making the network more efficient and capable of capturing important information. The process is repeated with another 2D convolutional layer with 64 filters and a ReLU activation function, followed by another MaxPooling layer.

A third 2D convolutional layer with 128 filters and ReLU activation is added, followed by another MaxPooling layer. The output of the third MaxPooling layer is then flattened into a one-dimensional vector. A fully connected (Dense) layer with 64 units and ReLU activation is added to further process the flattened features. Finally, there is an output Dense layer with a single unit and a sigmoid activation function ensures that the model’s prediction falls within the range [0, 1], allowing it to produce a probability indicating the likelihood of each input sample belonging to the positive class. This last layer is responsible for binary classification, as it outputs a probability value between 0 and 1.

The model is trained using 15 epochs and 32 batches, and it aims to learn patterns and features from the input data to perform the binary classification task.

## V. Results and Discussion

### A. Simulation Setup for Data Generation

We simulated three experiments (*N*_*sim*_ = 3) to generate training and test data. In each simulation, we considered a droplet of volume 10 *µ* L containing IPTG placed on a surface. The droplet was like a circle with a radius of 0.015 m and height of *h*_0_ = 1.4× 10^−5^ m.

For the first experiment, we imagined that the center of the droplet was right in the middle of the circle (*r* = 0 m), for the second experiment it was at the edge of the circle (*r* = 0.015 m), and for the third experiment, it was further away from the center (*r* = 0.03) m. The shape of the droplet was always like a circle.

Obviously, the system geometry is circular symmetric. Then, without loss of generality, we have assumed *θ* = 0 for all three release points. The number of molecules inside the tiny droplet was set at 10^4^, 10^5^, and 10^6^ for the first, second, and third simulations respectively. In other words, we are carrying out these simulations with different numbers of molecules in the droplet for the three experiments. This covers a wide range of stimulus concentrations. The diffusion coefficients in the agar and bacteria biofilm are considered as *D*_*a*_ = 10^−11^ m^2^ s^−1^ and *D*_*b*_ = 10^−8^ m^2^ s^−1^, respectively [22].

We have used a grid matrix with dimensions of 50 × 50 (with *N*_*x*_ = 50 and *N*_*y*_ = 50). Out of the total 2500 squares in this matrix, *N*_*s*_ = 1789 squares fall within the circular Petri dish. This means that we can gather 1789 data samples from each simulation, resulting in a total of 3 × 1789 samples. Table I provides the detailed structure of the proposed 2D-CNN.

**TABLE I:** Detailed Structure of 2D-CNN.

| Layer | Filter Size | Output Data Shape |
| --- | --- | --- |
| Input | - | (50, 50, 5) |
| Conv2D | 5×5 | (46, 46, 32) |
| MaxPooling2D | (2, 2) | (23, 23, 32) |
| Conv2D | 5×5 | (19, 19, 64) |
| MaxPooling2D | (2, 2) | (9, 9, 64) |
| Conv2D | 5×5 | (5, 5, 128) |
| MaxPooling2D | (2, 2) | (2, 2, 128) |
| Flatten | - | 512 |
| Dense | - | 64 |
| Dense | - | 1 |

In each simulation, when the system reaches a stable state, the bacteria colonies within each small section of the Petri dish may or may not produce GFP, i.e., the gene may be expressed or not. This depends on various factors, including the distance from release points and the boundary. Fig. 4 displays both the area where droplets are released and the labels for the square pieces within the Petri dish for all three simulations.

**Fig. 4:**
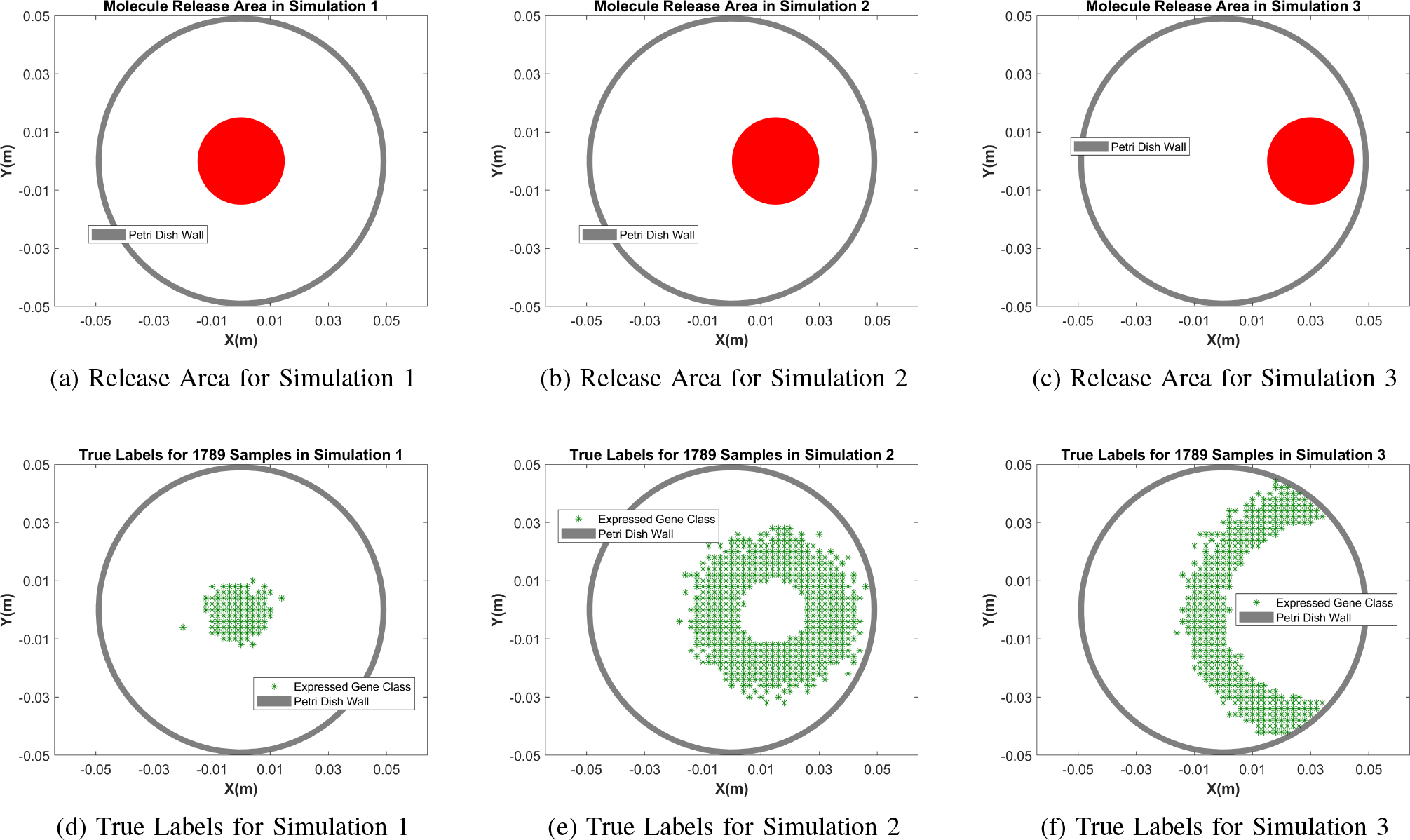
Released area and true labels for Simulations 1, 2, and 3 for generating 3 × 1789 data samples.

In the subfigures along the bottom half of Fig. 4, the green star symbols represent square areas where the genes were expressed and produced GFP. The plain white represents the pieces where the gene was not expressed (i.e., GFP was not produced). The gene expression corresponds to received IPTG concentrations between *C*_*L*_ = 10*/*(0.1*/*50)^2^ and *C*_*U*_ = 100*/*(0.1*/*50)^2^. The concentration of IPTG soaking into the release area is not uniform as described in Section III-A and is calculated using the model that takes into account the shrinking of the droplet, as described by equation (1). In this equation, the values are *k*_*s*_ = 1.3 × 10^−6^ ms^−1^ and *h*_*d*_ = 1.4 × 10^−5^ m.

Using this simulation configuration, we produced a total of 5367 data samples. Each input data has a structure resembling an image, with dimensions of 50 × 50 and 5 different channels. This setup is designed to work well with the proposed 2DCNN model.

Additionally, we generated a separate set of data for evaluation purposes, achieved through Simulation 4. In this simulation, we released a total of 500,000 molecules, and the droplet’s center was positioned at a distance of 0.02 m from the center (*r* = 0.02 m). In Fig. 5, the top part of the figure presents both the release area and the actual labels used for this particular simulation.

**Fig. 5:**
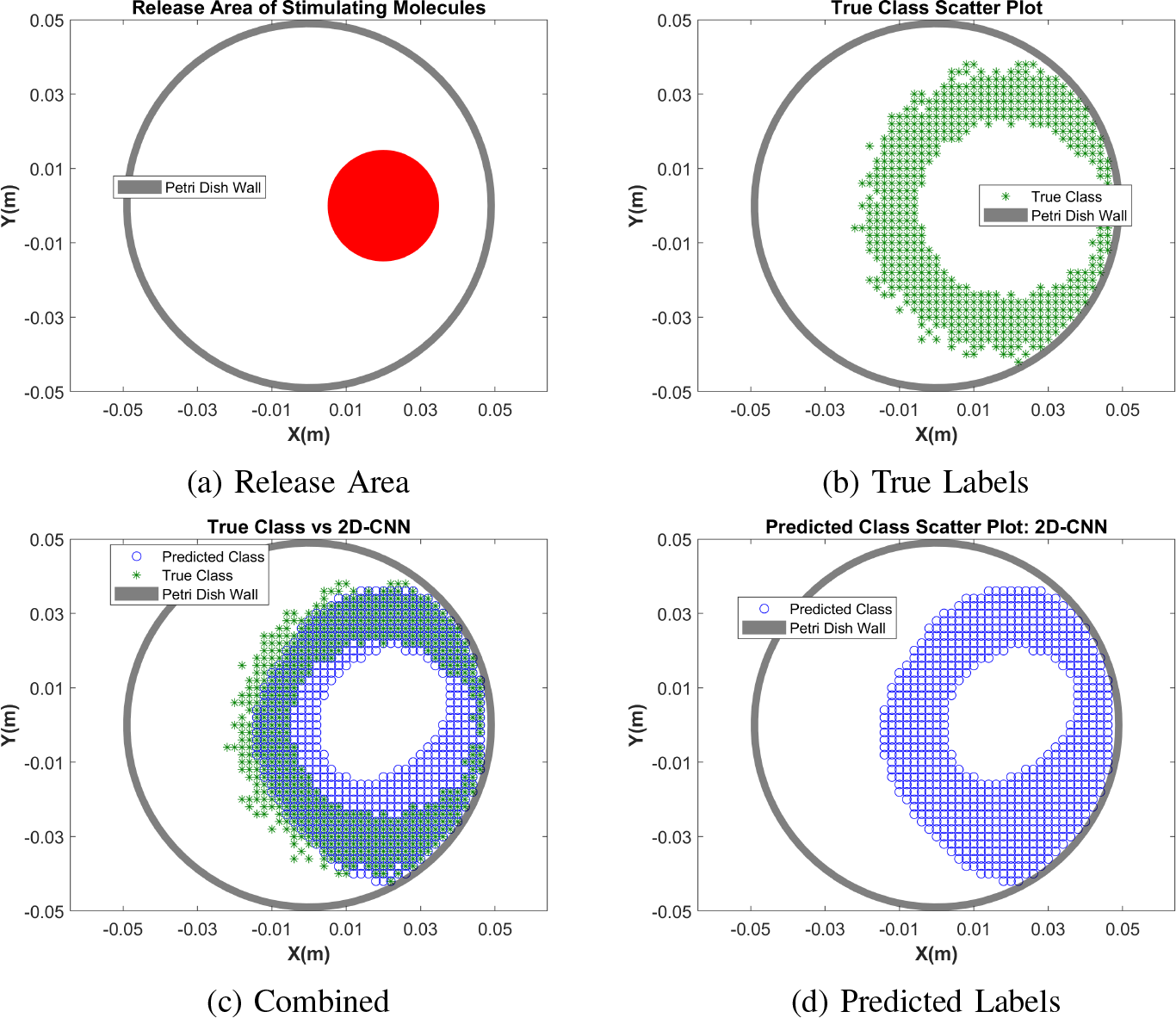
Visual evaluation of 2D CNN prediction of gene expression for test data (Simulation 4) that includes 1789 samples.

### B. Results

We used the data that we created through our simulation setup to train the proposed 2D-CNN model. This method was implemented using Python 3.10 along with the TensorFlow framework. For training, we used a loss function called categorical binary cross-entropy, and the training process was carried out using the Adam optimizer. We observe the general progress of the training in Fig. 6, which displays the training loss and accuracy for both the validation and test datasets.

**Fig. 6:**
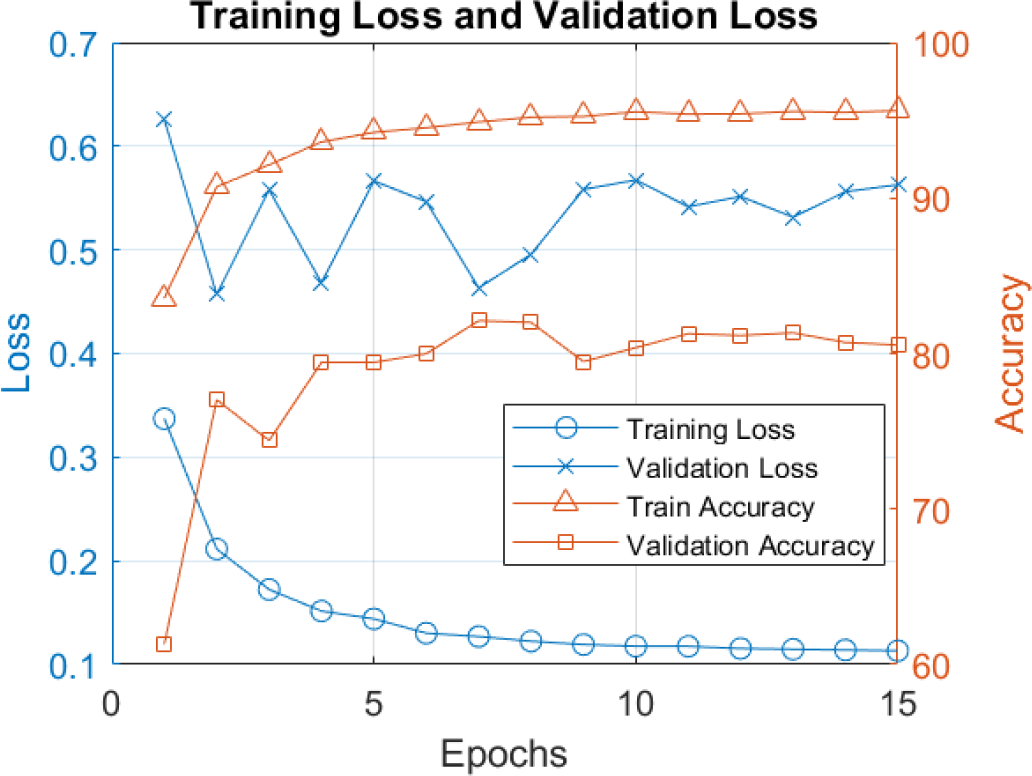
Training loss and accuracy of proposed 2D CNN model.

The proposed 2D-CNN model achieves an accuracy of 0.81 when evaluated on the test data. In Fig. 5 (bottom), the left side shows the labels predicted by the 2D-CNN, and on the right, a visual comparison is made between the actual labels and those predicted. It’s clear from the comparison that the 2D-CNN, utilizing the proposed data structure, has learned the gene expression system on the plate with high performance.

The confusion matrix is available in Table II. The corresponding false positive and false negative rates are both about 0.19 respectively, which is relatively low for this particular application considering the visual evaluation. Notably, when examining the combined scatter plot (c) in Fig. 5, we observe that the errors in predictions occur mostly at the border of the gene-expressed region. This phenomenon could be attributed to the inherent noise in diffusion across the plate, causing some uncertainty in delineating the precise borders of the expressed region. This is evident in the scatter plot of the true class in Fig. 5 upper right.

**TABLE II:** Confusion Matrix for 2D CNN.

|  |  | Predicted |  |
| --- | --- | --- | --- |
|  |  | Positive | Negative |
| Actual | Positive | 975 | 239 |
|  | Negative | 109 | 466 |

We conducted a comparison of the performance metrics of the 2D-CNN with several benchmark models including Linear SVM, Radial Basis Function SVM, Decision Tree, and Random Forest. In Fig. 7, we see the gene expression patterns predicted by all these benchmarks for the test data. These plots clearly illustrate that the 2D-CNN outperforms the benchmarks. For a detailed comparison of performance, Table III provides metrics including precision, recall, F1-score, and accuracy for both the CNN and the benchmark models.

**Fig. 7:**
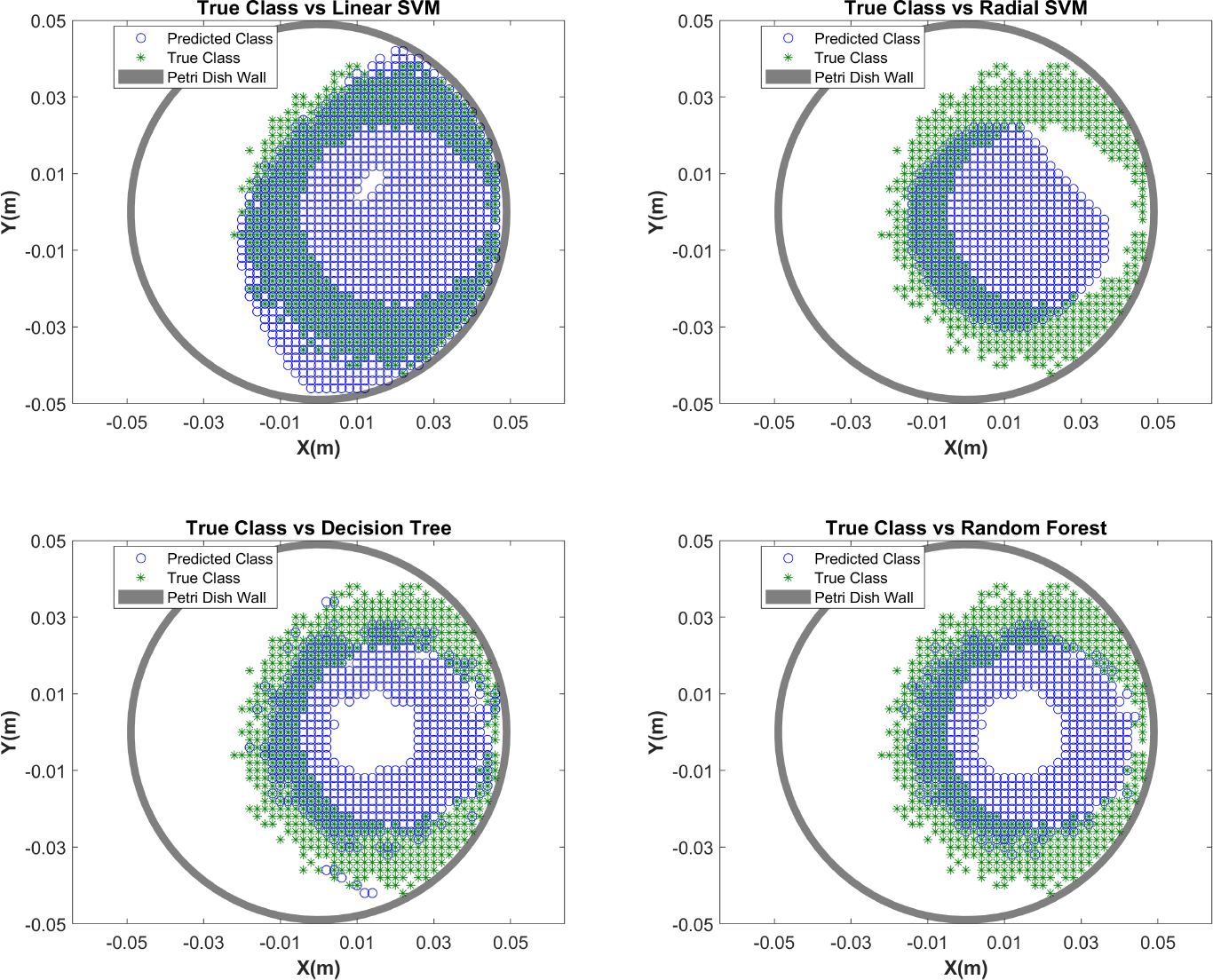
Gene expression prediction by benchmarks of Linear SVM, RBF SVM, Decision Tree, and Random Forest.

**TABLE III:** Comparison of Model Performance.

| Model | Precision | Recall | F1-Score | Accuracy |
| --- | --- | --- | --- | --- |
| 2D CNN | 0.68 | 0.81 | 0.73 | 0.81 |
| Linear SVM | 0.48 | 0.91 | 0.63 | 0.66 |
| RBF SVM | 0.33 | 0.30 | 0.31 | 0.58 |
| Decision Tree | 0.38 | 0.38 | 0.38 | 0.60 |
| Random Forest | 0.36 | 0.34 | 0.35 | 0.60 |

While the accuracy of Linear SVM is 0.66, the Linear SVM model exhibits a high recall value of 0.91, indicating a low rate of type II error (i.e., False Negative). The profile prediction of the Linear SVM, shown in Fig. 7 (Upper left), demonstrates that the hyperplane provided by this approach divides the plate into two continuous regions: the positive label area that corresponds to both the gene-expressed area producing GFP and also the area without gene expression due to an excessive concentration of IPTG which is labelled, while the other encompasses the remaining plate area labeled as negative. This strategy enables Linear SVM to correctly predict almost all positive labels. However, it struggles with predicting negative labels (non-expressed regions), leading to a high rate of type I error (False Positive) (1-0.48) and an overall accuracy of only 0.66. This discrepancy arises because the provided data structure for training doesn’t yield a hyperplane capable of accurately classifying elements in the non-expressed region due to the high concentration of molecules (surrounded by the expressed region). Similarly, the Radial SVM also faces difficulty distinguishing within the expressed region, where the IPTG concentration is high and genes are not expressed.

Decision Tree and Random Forest models (with 100 estimators) yield similar outcomes. In contrast to the Linear SVM, they exhibit improved performance in identifying the inner non-expressed region, as anticipated. However, they struggle to detect positive labels compared to the Linear SVM. As a result, Decision Tree and Random Forest both achieve an overall accuracy of 0.6.

## VI. Conclusion

In this paper, we introduced DeepGEEP, a deep learning approach aimed at predicting gene expression profiles based on the observed stimulus release pattern. DeepGEEP is able to reduce the resource and time-intensive experiments within Petri dishes that are very common across microbiology, biotechnology, and synthetic biology domains. Our proposed method incorporates a novel data generation approach that attains high accuracy with just a few experiments. Leveraging this training data structure, we designed a 2D-CNN model to predict gene expression patterns that attains 81% accuracy. We compared the proposed model against benchmark methods, including Linear SVM, Radial SVM, Decision Tree, and Random Forest.

Using simulations provided an adaptable, cost-effective means of gaining insights to refine our proposed DeepGEEP approach and as a proof of concept that suggest that the approach will be effective with just a limited number of physical lab experiments. Looking ahead, our focus will shift to the application of this structure on real experimental data. We also intend to design additional features that could effectively instruct the 2D-CNN to make more accurate predictions. Specifically, we aim to define composite features capable of encapsulating both diffusion and reaction properties inherent in the input data. Our current framework assumes a uniform spread of bacteria across the plate. Future efforts will involve introducing specified regions for bacterial activity, potentially leading to structural asymmetry. Furthermore, we aim to expand our approach to encompass variable release regions, moving beyond the current assumption of a constant release area. To accommodate all of these generalizations, we may need to embrace more intricate deep learning architectures, such as graph CNNs.

## Acknowledgment

This work was supported by the Engineering and Physical Sciences Research Council [grant number EP/V030493/1]. For the purpose of open access, the author has applied a Creative Commons Attribution (CC-BY) licence to any Author Accepted Manuscript version arising from this submission.

## Footnotes

1 ’cel” refers to the number of cells (bacteria)

